# Self-Supervised Transformer Model Training for a Sleep-EEG Foundation Model

**DOI:** 10.1101/2024.01.18.576245

**Authors:** Mattson Ogg, William G. Coon

## Abstract

The American Academy of Sleep Medicine (AASM) recognizes five sleep/wake states (Wake, N1, N2, N3, REM), yet this classification schema provides only a high-level summary of sleep and likely overlooks important neurological or health information. New, data-driven approaches are needed to more deeply probe the information content of sleep signals. Here we present a self-supervised approach that learns the structure embedded in large quantities of neurophysiological sleep data. This masked transformer training procedure is inspired by high performing self-supervised methods developed for speech transcription. We show that self-supervised pre-training matches or outperforms supervised sleep stage classification, especially when labeled data or compute-power is limited. Perhaps more importantly, we also show that our pre-trained model is flexible and can be fine-tuned to perform well on new EEG recording montages not seen in training, and for new tasks including distinguishing individuals or quantifying “brain age” (a potential health biomarker). This suggests that modern methods can automatically learn information that is potentially overlooked by the 5-class sleep staging schema, laying the groundwork for new sleep scoring schemas and further data-driven exploration of sleep.

## I. Introduction

Sleep is a fundamental biological process that promotes cognition [1] as well as mental [2] and physical health [3]. Sleep disturbances have also been linked to myriad neuropsychiatric conditions [4,5] that reduce quality of life for individuals and impose a large public health burden. The importance of sleep along with increases in the quality of consumer grade wearable sensors has generated excitement about health monitoring efforts that could flag changes for early clinical intervention or promote well-being by improving sleep habits [6].

Traditionally, sleep signals were analyzed by experts trained to assess signal features by eye. Each 30-second segment received one of five labels corresponding to either wakefulness (W), one of three non-rapid eye movement (NREM) sleep stages (N1, N2, N3), or rapid eye movement (REM) sleep [7]. Five-class sleep scoring is still the backbone of sleep medicine today, but carrying out this process manually is laborious and creates a bottleneck for large scale efforts. A pivot to the use of automated techniques has been accelerated by the application of deep learning to large quantities of public sleep data (see [8] for review). Today, automated sleep scoring algorithms perform consistently as well as human experts [9-12], whose maximum inter-rater agreement is 80-82% [13,14].

Yet, while AI and data-driven approaches have generated numerous breakthroughs, they will not realize their full potential if confined to traditional five-stage sleep labels. For example, a focus on sleep stages is subject to the shortcomings of traditional sleep scoring including ambiguity in label application (see 80-82% reliability above [13,14]). Other issues include a “one-size-fits-all” lack of personalization, and concessions to efficiency aimed at reducing technician workload (e.g., labeling 30-second windows is less time-consuming than labeling 10-second windows).

Taken together, it is likely that supervised methods overlook important neurological or health information not captured by the standard sleep labeling task [15,16] that could address some of the shortcomings that still exist in sleep medicine. For example, micro-scale neurophysiological artifacts observed during sleep are related to health outcomes that might not relate directly to sleep stages [17,18].

An alternative approach that could support the extraction of additional information from sleep signals is the use of self-supervised neural network training methods to learn the structure that exists in sleep data. Such hypothesis-free methods aim to learn generalizable representations that map the latent information space that exists within a (typically very large) set of data and could distill a variety of clinically relevant patterns (e.g., sleep stages, apnea classifications, disease states, etc.). Deploying these techniques on the huge quantities of publicly available sleep data (and on electroencephalographic, or EEG, signals in particular, which encode most of the important sleep landmarks as well as other health information [19]) could unlock further advances in sleep medicine and better leverage available data resources.

In this work we report an initial investigation into self-supervised learning methods applied to EEG signals from a large corpus of public polysomnography (PSG) data. Critically, pre-training these methods does not require PSG annotations or sleep labels. To do this we adapted an approach developed for self-supervised transformer model training in support of speech recognition (“HuBERT” [20]). In other work we have shown that transformer models are adept at scoring sleep [21]. Here, we broaden that work to show that these models are also adept at self-supervised learning and extract nuanced features of sleep from EEG beyond canonical sleep-stage labels.

## II. Methods

### A. Pre-Training Data and Processing

Self-supervised learning requires a large corpus of unlabeled data for pre-training. To meet this demand, we assembled a dataset of 10,897 sleep sessions from 9,013 individuals from public sources hosted on the National Sleep Research Resource [22] (NSRR) for pre-training: the Multi-Ethnic Study of Atherosclerosis [23] (1461 sessions from 1461 individuals aged 54 to 90, 53.2% female), the MrOS Sleep Study [24] (2327 sessions from 1934 individuals aged 67 to 90, 0% female), the Sleep Heart Health Study [25] (4197 sessions from 3461 individuals aged 39 to 90, 51.7% female), the Wisconsin Sleep Cohort [26] (1658 sessions from 920 individuals aged 37 to 78, 46.5% female), the Nationwide Children’s Hospital Sleep DataBank [27] (197 sessions from 180 individuals aged 18 to 58, 50.8% female), the Study of Osteopathic Fractures [28] (398 sessions from 398 individuals aged 76 to 90, 100% female), and the Cleveland Family Study [29] (659 sessions from 659 individuals aged 7 to 89, 55.1% female). Data from an additional 915 individuals (one night each) from these datasets was held out for validation.

From each sleep session we retained a central EEG channel (C3 or C4, similar to [10]). Each EEG channel was resampled to a 100Hz sampling rate, normalized via median subtraction, scaled to achieve an inter quartile range (IQR) of unity (1.0) and truncated to remain within +/-20 IQR. Data were then segmented into 30-second epochs based on the sleep-stage annotations, ignoring epochs not explicitly assigned a sleep-stage (i.e., removing any artifactual or unscored segments). These 30-second epochs were then concatenated consecutively into 101-epoch sequences every 25 epochs (i.e., oversampling the data by a factor of four for pre-training).

### B. Self-Supervised Learning

Our self-supervised method trains a transformer model to recognize sequences of simple k-means labels that are discovered within the training data, an approach based on HuBERT [20]. See Figure 1 for a schematic of our approach. These k-means labels are discovered a-priori by converting each 101-epoch sequence of EEG time series data to a spectrogram of 30-second time bins (based on the YASA [12] algorithm), and then deriving a k-means label based on the frequency information in each time bin. These k-means labels are then used instead of sleep stage labels during pre-training.

**Fig. 1.**
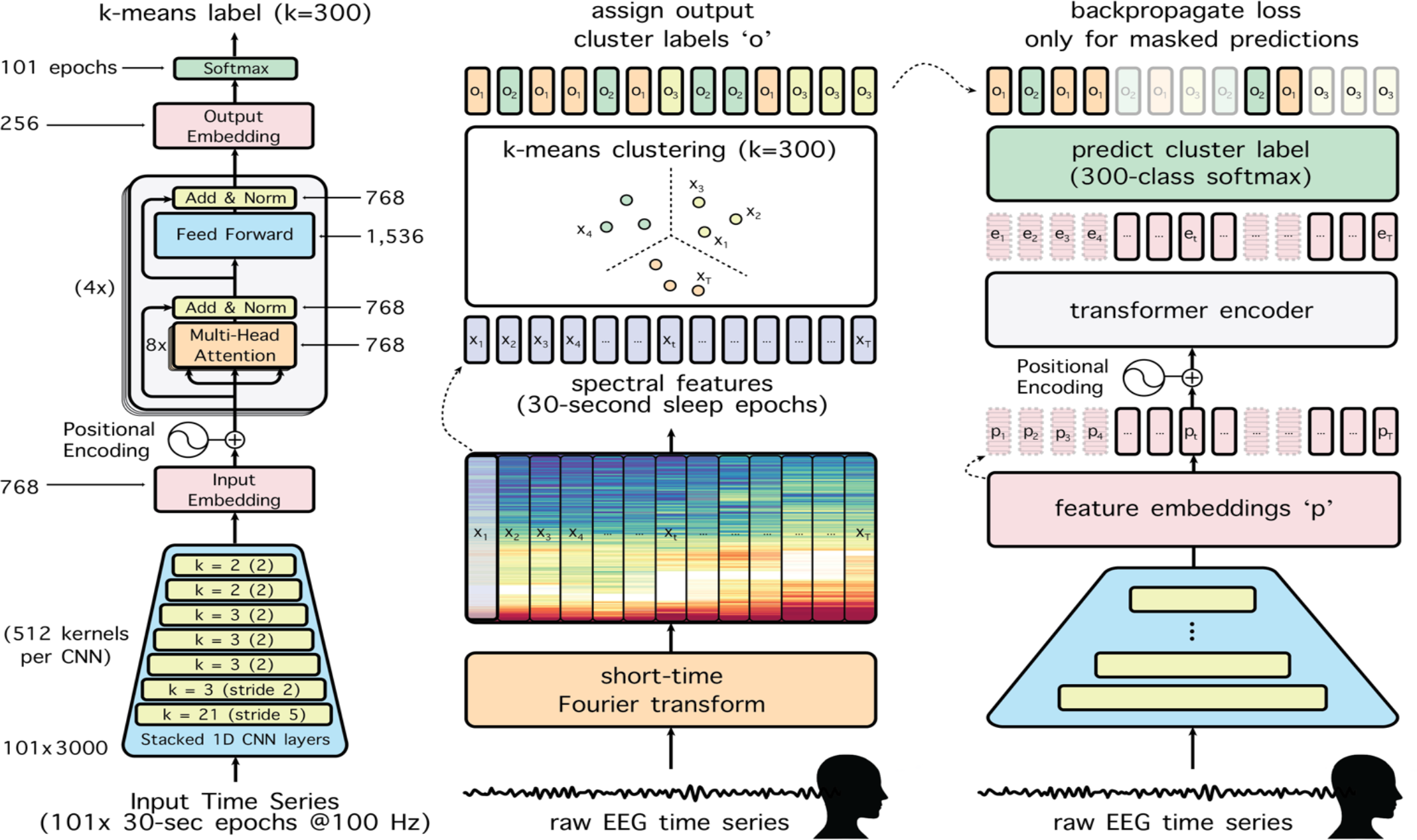
Schematic of our transformer model and self-supervised training approach. *The model is trained to map input time series data to a masked sequence of k-means labels discovered within the data. The model (left) comprises a 1D-CNN frontend, linear projection (p_n_), transformer, a second linear projection (e_n_) and an output layer (o_n_). K-means labels are derived from a spectrogram representation of the EEG of the training data (middle). During training (right) only performance within masked epochs (bolded in the output layer) contributes to loss. After training, for some fine-tuning experiments (participant recognition and age prediction), the layers e and o are discarded and the transformer output is averaged to distill an embedding for fine-tuning.*

We initially derived a 300-cluster k-means solution, splitting the difference between HuBERT’s original [20] 100-then-500-cluster training phases to accommodate one phase of self-supervised training. We obtained this solution using one randomly selected 101-epoch segment from each participant in the training data (using the mini-batch k-means algorithm). We then applied this solution (i.e., generating one of the 300 k-mean labels for each 30-second epoch in each 101-epoch sequence) to the training and validation data. We include two additional (dummy) label classes to capture the very small number of k-means failures (which arise from infrequent inf or NaN values in the spectrogram, respectively).

During training we mask (with zeros) portions of the input corresponding regions of the 101-epoch k-means sequences before sending them to the transformer. The loss function captures how well the model correctly assigns the appropriate sequence of k-means labels to masked portions of the input data. The input data are masked in blocks of ten epochs where each of the 101-epochs in the input sequence has an 8% probability of being the starting point of a masked block.

Our transformer model first uses a series of seven 1-dimensional convolutional layers (each with 512 channels, strides = 5,2,2,2,2,2,2 and kernel widths = 21,3,3,3,3,2,2 across layers) that learn filter kernels to optimally process the data based on the input time series. Each of these layers involves a 1-dimensional convolution, layer-norm and a gelu activation function. The output of these convolutions then undergoes a linear projection from 512 to 768 units with gelu activation followed by positional encoding and then four transformer encoder layers (with 8 attention heads, an inner feed forward dimension of 1536, gelu activation and 5% layer drop probability). The output of the transformer undergoes 1-dimensional adaptive average pooling to align with the label sequence of length 101. This averaged transformer output is projected to a 256-dimensional embedding followed by gelu activation and then the output layer. Masking is applied after the convolutional front end, prior to positional encoding and the transformer. The full model contains 28.5 million trainable parameters (occupying 129.5 MB on disk).

We trained our transformer model for 40 epochs (i.e., 40 passes through the entire pre-training corpus) using a batch size of 32 sequences. The model was trained to minimize cross-entropy on the k-means labels for the masked portions of the input sequences. We used the Adam optimizer with a linear ramp up learning rate scheduler (0.00001 to 0.000025 for the first 15 epochs). At the end of pre-training we retained the model from the epoch with the lowest validation loss.

Note that the data, preprocessing and model architecture for this study are identical to those in our related report examining transformers for supervised sleep scoring [21].

### C. Fine-Tuning Experiments

One major challenge in machine learning is data paucity. Limited data (or labels) may not be of sufficient volume to support training a large, high-performing supervised model from scratch without overfitting. Fine-tuning can be used in these cases by first pre-training a model on a large volume of data, or a more general task, to learn useful features before a second stage of training where the model is briefly trained on a smaller quantity of (usually more in-domain) data or labels. Transfer learning is a similar procedure (which also uses fine-tuning), where a second-stage training run maps the model (or model embeddings) to another dataset where the task or output is more dissimilar from pre-training. Fine-tuning pre-trained models for speech transcription has been extremely effective when labelled data are limited [20].

We carried out two sets of experiments to understand if our model delivered benefits similar to foundation models in other domains. Specifically, 1) does our model make efficient use of labeled data?, and 2) can our model be flexibly fine-tuned to support diverse downstream tasks? We evaluated our fine-tuning experiments on the validation data and Sleep-EDF corpus. The Sleep-EDF data was preprocessed in the same manner as the NSRR data, only using Fpz-Cz as the EEG channel and no oversampling for evaluation folds.

The first set of experiments assessed our model’s performance when fine-tuned for a standard sleep stage sequence learning task given limited labeled data. We fine-tuned our model to classify the sleep stages (Wake, N1, N2, N3, REM) of each epoch in our 101-epoch sequences using annotated sleep data from 10, 100, 500 or 1000 individuals (one night each) or the entire training corpus. Models were then evaluated using the Sleep-EDF data. We used two baseline models against which we could gauge the performance of our self-supervised model. Both baseline models used the same transformer architecture, training data and hyperparameters as our self-supervised model, and differed only in their use/non-use of k-means pseudo-labels and/or masking. One baseline transformer model underwent traditional supervised training to perform the standard five-way sleep stage classification task (i.e., a supervised model trained from scratch on each limited dataset). The other baseline model was pre-trained to classify the same sequence of k-means labels we used for self-supervised training but without any masking during pre-training. This latter model allowed us to assess how much of the performance of our full self-supervised model was due to the k-means labels alone or to the masking procedure during pre-training. Models were run for 50 epochs with the same Adam optimizer. We allowed the pre-trained models to update all weights during fine-tuning for sleep stage classification.

The second set of experiments sought to understand what other, non-sleep stage information the model encoded during pre-training. For this we focused on participant recognition and age prediction. The participant recognition and age prediction experiments were carried out on the Sleep-EDF dataset. We used split-half cross-validation (folds: night-1/night-2 for subject recognition, and folds: even-subject-ID/odd-subject-ID for age prediction) and we report performance averaged over both folds. For these experiments we froze the weights of all models (the full self-supervised model, the model trained with k-means labels but no masking, and supervised five-way sleep stage classification model trained on the full training corpus). Specifically, for each 101-epoch example sequence in the Sleep-EDF dataset, we averaged the output of the last (frozen) transformer layer from each model, resulting in one 768-dimensional embedding for each model for each sequence. The training folds for each experiment used the four-times oversampled Sleep-EDF sequences while the evaluation folds always used non-overlapping sequences. In these experiments training can be described as learning a simple “linear read-out” layer between the average pre-trained transformer output (embedding) and output units for each task. The subject recognition experiments involved one output unit per participant ID (trained with cross entropy loss) and the age prediction experiments used a single output unit to predict age (trained with MSE loss). These simple feed-forward (linear read-out) models were trained for 50 epochs using stochastic gradient descent (batch-size of 64, learning rate of 0.001 which was reduced if there was no performance improvement for 3 epochs). For these experiments we also trained another transformer model (same architecture) from scratch on each subject ID or age prediction task as an additional baseline (with the same hyper-parameters as the sequence classification models above, and a batch size of 32).

## III. Results

Our model was able to learn the pre-training task effectively. Performance at recognizing k-means labels within masked portions of the validation data peaked at a top-1 accuracy of 36.59% and a top-5 accuracy of 76.05% after 250k training steps (epoch 36). However, classification of masked k-means labels is (by design) a somewhat arbitrary task, so to understand how effective our self-supervised pre-training strategy was, we submitted this pre-trained model to different fine-tuning experiments.

### A. Fine-Tuning for Sleep-Stage Classification with Limited Data

As shown in Table 1, our self-supervised transformer model that predicted masked sequences of k-means labels outperformed the baseline models when data was limited, and also converged faster (i.e. requiring fewer training epochs) than both supervised models trained on limited data and a model pre-trained to predict k-means labels but without masking. We note that when using annotated data from 500 or more participants, sleep stage classification performance began to converge among models, but our self-supervised model often still converged in fewer epochs than the other models. All models performed exceptionally well on the internal validation data but our self-supervised model generalized its performance to perform well on the Sleep-EDF dataset better than the other models, especially when using smaller labeled datasets. The lower fine-tuning performance of the baseline model pre-trained to predict sequences of k-means labels (but trained without sequence masking) indicates that the high performance of our full self-supervised model was enabled by both the k-means labels in pre-training and the masked prediction task for portions of the input sequences.

**TABLE I.** Fine-Tuning for Sleep-Stage Classification Task.

| Pre-train Target | Pre-Train Task | Finetuning | Labeled Sleep Sessions | Internal Val. (Accuracy %) | Best Val. Epoch | Sleep-EDF (Accuracy %) |
| --- | --- | --- | --- | --- | --- | --- |
| K-means | Predict Masked | Fine-tune Pre-Trained | 10 | 75.82 | 9 | 75.4 |
| K-means | Predict Masked | Fine-tune Pre-Trained | 100 | 81.76 | 5 | 76 |
| K-means | Predict Masked | Fine-tune Pre-Trained | 500 | 84.11 | 24 | 77.85 |
| K-means | Predict Masked | Fine-tune Pre-Trained | 1000 | 84.82 | 28 | 76.98 |
| K-means | Predict Masked | Fine-tune Pre-Trained | All | 86.54 | 46 | 77.3 |
| K-means | Seq. Prediction | Fine-tune Pre-Trained | 10 | 66.68 | 8 | 62.86 |
| K-means | Seq. Prediction | Fine-tune Pre-Trained | 100 | 80.04 | 17 | 75.5 |
| K-means | Seq. Prediction | Fine-tune Pre-Trained | 500 | 83.54 | 21 | 75.13 |
| K-means | Seq. Prediction | Fine-tune Pre-Trained | 1000 | 85.03 | 34 | 78.39 |
| K-means | Seq. Prediction | Fine-tune Pre-Trained | All | 86.62 | 49 | 76.51 |
| Sleep stages | Seq. Prediction | Trained from Scratch | 10 | 69.75 | 40 | 61.92 |
| Sleep stages | Seq. Prediction | Trained from Scratch | 100 | 80.2 | 40 | 70.9 |
| Sleep stages | Seq. Prediction | Trained from Scratch | 500 | 84.04 | 43 | 77.88 |
| Sleep stages | Seq. Prediction | Trained from Scratch | 1000 | 84.77 | 30 | 74.5 |
| Sleep stages | Seq. Prediction | Trained from Scratch | All | 86.59 | 49 | 77.41 |

### B. Fine-Tuning for Non-Sleep Staging Tasks: Individual Identification, Age Prediction

Beyond predicting sleep stage labels, the second set of experiments demonstrates that our self-supervised model performs well on diverse downstream tasks (Table 2). Our self-supervised model’s representations can support recognition of over 70 different participants across different nights of sleep, and at higher accuracy than a model pre-trained for sleep-stage classification, a model pre-trained on k-means sequences but without masking or a trained from scratch directly for this task.

**TABLE II.** Fine-Tuning for Non-Sleep Staging Tasks (Participant Recognition AND Age Prediction)

| Pre-train Target | Pre-train Task | Finetuning | ID Top-1 | ID Top-5 | Age Pearson r | Age MAE |
| --- | --- | --- | --- | --- | --- | --- |
|  |  |  | Accuracy (%) | Accuracy (%) |  |  |
| K-means | Predict Masked | Fine-tune Embed | 55.94 | 75.045 | 0.677 | 13.38 |
| K-means | Seq. Prediction | Fine-tune Embed | 24.72 | 48.405 | 0.3607 | 17.775 |
| Sleep stages | Seq. Prediction | Fine-tune Embed | 17.305 | 41.75 | 0.4101 | 18.63 |
|  |  | From Scratch | 1.1465 | 7.09 | 0.346 | 16.145 |

Our self-supervised model was also able to support accurate age prediction within plus or minus 15 years (mean absolute error). Recent work suggests that this could be useful as a potential health biomarker [30]. The performance of our model approaches that of studies using fMRI data [31] although some purpose-built EEG models (trained for age prediction with much more data) perform better [18]. MRI is expensive so the ability to estimate brain age with inexpensive EEG, which can also be used in the home, provides an attractive alternative.

## IV. Discussion

Together our results indicate that foundation models can deliver for brain data (here, sleep EEG signals) many of the same benefits that foundation models have provided in other domains like text and speech. Our self-supervised pre-trained transformer model is efficient (obtaining high performance with limited labeled data) and flexible (performing well when fine-tuned on diverse downstream tasks). It will be interesting to explore how additional sources of data or objectives in pretraining could influence performance, and to evaluate the model on more diverse downstream tasks.

We also note some limitations, notably that currently the model only uses EEG, while additional sleep and health information [19] could be gleaned from additional signals. Evaluations on more diverse data and input signals are needed, which we are beginning to address in ongoing work. Also, evaluation conditions not seen during training such as different clinical populations, recording montages, or signal quality could hurt performance (sometimes called a domain shift) if these characteristics are not integrated into pre-training or fine-tuning. For example, we were able to perform well on the different (bi-polar referenced) montage in the Sleep-EDF data following fine-tuning. We also expect performance to be improved following further optimizations to the clustering solution, model architecture and other hyperparameters.

We are keenly interested in using this approach to identify specific clinical states as well as recognizing aberrant sleep patterns or clinically relevant neural activity during sleep. This could pave the way for more information-rich, data-driven sleep state analysis schemas that will aid in diagnosis or facilitate the interpretation of sleep patterns to quantify and predict health or disease outcomes. In this way, interpreting sleep signals with the aid of foundation models and their derivatives could be truly transformative for the practice of sleep medicine.

## Acknowledgment

The authors acknowledge the support from the Independent Research and Development (IRAD) Fund from the Research and Exploratory Development Mission Area of the Johns Hopkins University Applied Physics Laboratory. The Cleveland Family Study (CFS) was supported by grants from the National Institutes of Health (HL46380, M01 RR00080-39, T32-HL07567, RO1-46380). The National Heart, Lung, and Blood Institute provided funding for the ancillary MrOS Sleep Study, “Outcomes of Sleep Disorders in Older Men,” under the following grant numbers: R01 HL071194, R01 HL070848, R01 HL070847, R01 HL070842, R01 HL070841, R01 HL070837, R01 HL070838, and R01 HL070839. The Multi-Ethnic Study of Atherosclerosis (MESA) Sleep Ancillary study was funded by NIH-NHLBI Association of Sleep Disorders with Cardiovascular Health Across Ethnic Groups (RO1 HL098433). MESA is supported by NHLBI funded contracts HHSN268201500003I, N01-HC-95159, N01-HC-95160, N01-HC-95161, N01-HC-95162, N01-HC-95163, N01-HC-95164, N01-HC-95165, N01-HC-95166, N01-HC-95167, N01-HC-95168 and N01-HC-95169 from the National Heart, Lung, and Blood Institute, and by cooperative agreements UL1-TR-000040, UL1-TR-001079, and UL1-TR-001420 funded by NCATS. NCH Sleep DataBank was supported by the National Institute of Biomedical Imaging and Bioengineering of the National Institutes of Health under Award Number R01EB025018. The Sleep Heart Health Study (SHHS) was supported by National Heart, Lung, and Blood Institute cooperative agreements U01HL53916 (University of California, Davis), U01HL53931 (New York University), U01HL53934 (University of Minnesota), U01HL53937 and U01HL64360 (Johns Hopkins University), U01HL53938 (University of Arizona), U01HL53940 (University of Washington), U01HL53941 (Boston University), and U01HL63463 (Case Western Reserve University). The Study of Osteoporotic Fractures (SOF) was supported by National Institutes of Health grants (AG021918, AG026720, AG05394, AG05407, AG08415, AR35582, AR35583, AR35584, RO1 AG005407, R01 AG027576-22, 2 R01 AG005394-22A1, 2 RO1 AG027574-22A1, HL40489, T32 AG000212-14). This Wisconsin Sleep Cohort Study was supported by the U.S. National Institutes of Health, National Heart, Lung, and Blood Institute (R01HL62252), National Institute on Aging (R01AG036838, R01AG058680), and the National Center for Research Resources (1UL1RR025011). The National Sleep Research Resource was supported by the U.S. National Institutes of Health, National Heart Lung and Blood Institute (R24 HL114473, 75N92019R002).

